# Specialization of actin isoforms derived from the loss of key interactions with regulatory factors

**DOI:** 10.1101/2021.02.09.430555

**Authors:** Micaela Boiero Sanders, Christopher P. Toret, Adrien Antkowiak, Robert C. Robinson, Alphée Michelot

## Abstract

A paradox of eukaryotic cells is that while some species assemble a complex actin cytoskeleton from a single ortholog, other species utilize a greater diversity of actin isoforms. The physiological consequences of using different actin isoforms, and the molecular mechanisms by which highly conserved actin isoforms are segregated into distinct networks, are poorly known. Here, we sought to understand how a simple biological system, composed of a unique actin and a limited set of actin-binding proteins, reacts to a switch to heterologous actin expression. Using yeast as a model system and biomimetic assays, we show that such perturbation causes drastic reorganization of the actin cytoskeleton. Our results indicate that defective interaction of a heterologous actin for important regulators of actin assembly limits certain actin assembly pathways while reinforcing others. Expression of two heterologous actin variants, each specialized in assembling a different network, rescues cytoskeletal organization and confers resistance to external perturbation. Hence, while species using a unique actin have homeostatic actin networks, actin assembly pathways in species using several actin isoforms may act more independently.

## Introduction

A fundamental characteristic of eukaryotic cells is the existence of an organized actin cytoskeleton. Dynamic actin filaments are assembled into diverse architectures which co-exist within one cytoplasm, each of which is involved in the exertion of forces for various cellular functions (Blanchoin *et al*, 2014). Key partners are families of actin-binding proteins (ABPs), which interact with actin monomers and filaments to regulate cytoskeletal organization and dynamics (Moseley & Goode, 2006; Pollard, 2016). Actin sequence is highly conserved across most eukaryotes, but while some cell types only express a single actin (for example yeasts), other cell types can express several similar actin isoforms (for example non-muscle mammalian cells express beta- and gamma-actins which are 99% identical), or even very different actin isoforms (for example, *Chlamydomonas reinhardtii* expresses two actins, IDA5 and NAP1, which are only 65% identical) (Gunning *et al*, 2015; Boiero Sanders *et al*, 2020). An extreme case is plants, which can express a multitude of actin isoforms (for example, *Zea mays* and *Arabidopsis thaliana* express 21 and 8 actin isoforms, respectively). Adding to this complexity, some actins can undergo partial post-translational modifications (PTMs), such as arginylation or acetylation, which modify their biochemical properties (A *et al*, 2020; Kashina, 2014; Boiero Sanders *et al*, 2020).

Hence, while a number of organisms are able to assemble a complex actin cytoskeleton from one (or a limited number) of actin isoforms, other organisms require the presence of multiple actin isoforms to generate such variability. In line with this idea, segregation of actin isoforms is observed *in vivo*. Results from different mammalian cell lines have found that beta-actin was located mainly in the contractile ring, stress fibers, filopodia and cell-cell contacts while gamma-actin was localized primarily in the cortex and lamellipodia (Dugina *et al*, 2009; Chen *et al*, 2017). In *Arabidopsis thaliana*, the main vegetative actin isoforms organize into different structures in epidermal cells (Kijima *et al*, 2018). However, it should be noted that expression in mice of a beta-coded gamma-actin, where the nucleotide sequence of beta-actin is modified minimally to express gamma-actin, led to viable mice with no detectable change in behavior (Vedula *et al*, 2017). This result indicates that at least in some cases, the absence of an actin isoform can be compensated by the expression of a similar isoform.

A particular challenge for the field is to understand how small differences at the molecular level lead to a major segregation of actin isoforms at the cellular level. To decipher the underlying mechanisms, it is natural to postulate that actin isoforms bear small yet significant biochemical differences. Our knowledge of the distinctions between actins is limited to a small number of actin orthologs (mainly *S. cerevisiae* Act1p, rabbit muscle actin, to a lesser extent beta- and gamma-actins, *S. pombe* Act1p and plant actins). Nonetheless, these studies reveal notable differences in their biochemical properties (Nefsky & Bretscher, 1992; Kim *et al*, 1996; Buzan & Frieden, 1996; Bryan & Rubenstein, 2005; Takaine & Mabuchi, 2007; Kijima *et al*, 2016), in their mechanical properties (Orlova *et al*, 2001; McCullough *et al*, 2011), and their ability to interact with the different actin-binding proteins (Nefsky & Bretscher, 1992; Eads *et al*, 1998; Takaine & Mabuchi, 2007; Ezezika *et al*, 2009; McCullough *et al*, 2011; Kang *et al*, 2014; Kijima *et al*, 2016), including nucleation factors of actin assembly (Ti & Pollard, 2011; Chen *et al*, 2017). How such differences account for spatial segregation of actin isoforms on a cellular scale remains unclear.

In this work, we investigated, from a general perspective, the molecular principles by which actin isoforms can be addressed to different networks. Analysis in a model system, that exploits at least two actins to perform various actin functions, would explain a particular mechanism in a relevant physiological context. However, the importance of actin renders genetic manipulations difficult, and the inter-connection of actin networks in such models complicates cellular analysis. Mammalian systems in particular express many ABP isoforms, which makes interpretation of molecular mechanisms combinatorially challenging. Furthermore, co-expression of multiple actin isoforms makes endogenous purification as a single species difficult, although new powerful protocols have been developed in recent years for their expression and purification (Hatano *et al*, 2018, 2020). To overcome these limitations, we decided to adopt an alternative strategy, by determining the consequences of heterologous actin expression in a system normally using a single actin. With this approach, we aimed at measuring the consequences of a perturbation caused by the use of a different actin at the level of the cell and its cytoskeleton. We decided to use the well-studied organism, budding yeast, for the simplicity of its genetics. Another advantage of budding yeast is that actin assembles predominantly into two well-defined structures. These structures are actin patches, which are sites of endocytosis and where actin filaments are short and branched by the Arp2/3 complex, and actin cables, which are central for maintenance of cell polarity and intra-cellular trafficking, and where actin filaments are nucleated by the formin family of proteins (Moseley & Goode, 2006). Lastly, budding yeast allows for clean purification of ABPs in a defined organismal context.

Our results demonstrate that actin functions are regulated both at the nucleotide level where defects in actin expression leads to cell growth defects, and at the amino acid level where expression of heterologous actins induce a massive reorganization of the actin cytoskeleton. We demonstrate that actin isoforms are used with different efficiencies by the distinct actin assembly pathways, resulting in their targeting to particular actin structures. Finally, dissection of the underlying molecular mechanisms allows us to propose an explanation of our results, and a general model of the molecular mechanisms enabling segregation of actin isoforms in cells.

## Results

### Generation of a library of yeast strains expressing a variety of actin orthologs

We created a library of *S. cerevisiae* strains that express different actin orthologs to evaluate the consequences of actin variation on yeast actin cytoskeleton assembly. In order to ensure that defects were not due to potential misfolding or non-functional actin, we selected a diversity of actins from other species rather than using directed mutations. This approach guarantees that the actin orthologs are functional in a biologically-relevant context, and maintain key physiological properties such as polymerization, depolymerization, nucleotide binding and hydrolysis.

We chose 122 different actins from species covering the entire eukaryotic phylogenetic tree for analysis (Table S1 and Fig. S1 A). We also computationally predicted ancestral sequences to extend the range of actin variant possibilities. Because the actin protein sequence is highly conserved across species, ancestral sequence reconstructions score with high confidence (Fig. S1 B). We obtained in total 223 actin sequences (including 101 ancestral actins), from which we selected 15 for analysis. These actin orthologs were chosen to cover a spectrum from the most similar to wild-type *S. cerevisiae*’s actin (Act1p) to very divergent actin orthologs, which represent a wide range of identities (from 99 to 84%) (Fig. 1A and S1 B-C, Table 1), and to display differences across all domains of the actin fold (Fig. 1 B-C and S1C).

**Figure 1.**
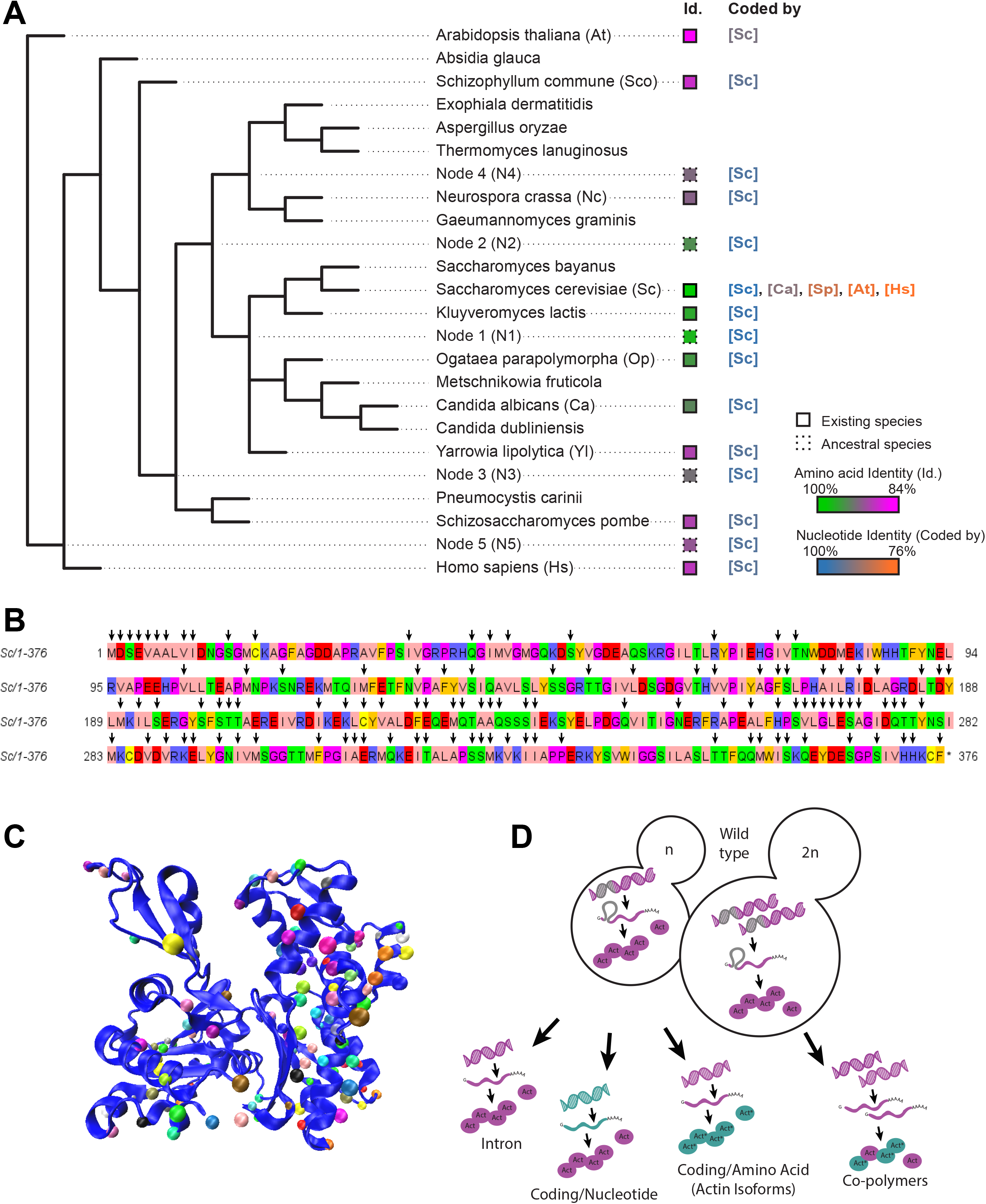
Variety of actins selected for this study and analysis strategies. **(A)** Simplified phylogenetic tree showing mainly the Dikarya subkingdom and including the external branches *Homo sapiens* (Hs) and *Arabidopsis thaliana* (At). The Id. column indicates amino acid sequences percentage identities, ranging from 100% (green) to 84% (magenta) identity to *S. cerevisiae*’s actin. Squares outlines are solid or dotted for sequences deriving from existing species or ancestral reconstruction, respectively. The “coded by” column indicates which coding sequences were originally used to code genes of interest. Nucleotide sequence identities are ranging from 100% (blue) to 76% (orange) compared to *S. cerevisiae*’s actin coding sequence. **(B)** Amino acid sequence of *Saccharomyces cerevisiae* actin. Arrows denote all the positions that are mutated in at least one of the actin variants tested in this study. **(C)** Schematic representation of *S. cerevisiae* actin 3D structure (1YAG (Vorobiev *et al*, 2003)), showing that mutations cover all regions of the protein. Dots indicate where mutations are located, using a different color code for all actins studied here. **(D)** Schematic showing the mutagenesis strategies applied in this study, enabling to question respectively the importance of actin’s intron, the nucleotide sequence, the amino acid sequence, and the effect of expressing copolymers. Green color indicates whether modifications are brought in the coding sequence (leading to expression of wild-type Act1 protein (pink) or in the amino acid sequence (leading to expression of an Act1* actin ortholog).

**Table 1.**
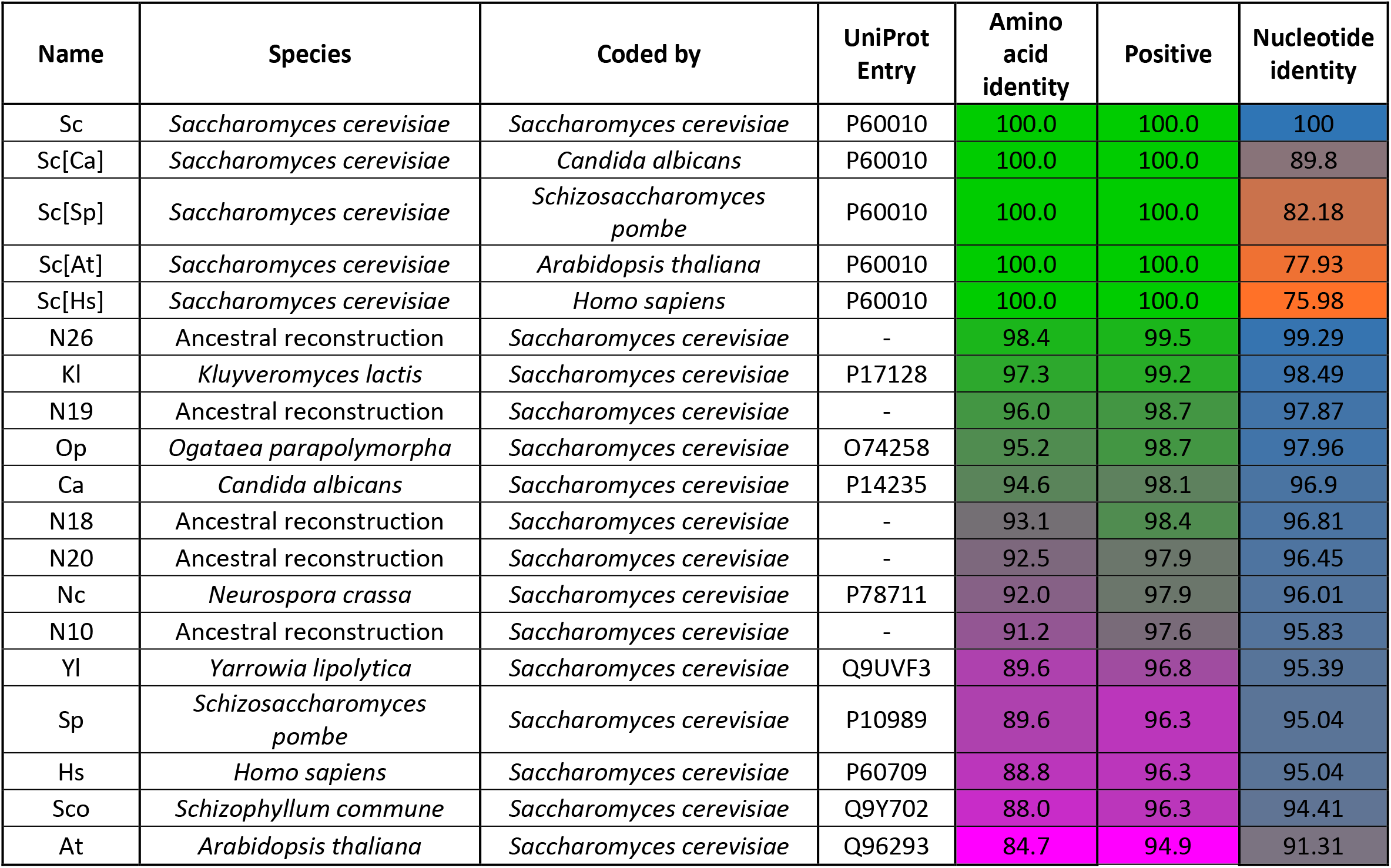
Details of actin variants. This table contains the information corresponding to all actins used in this study. Name corresponds to the names used in the manuscript. Species corresponds to species where the amino acid sequence is normally expressed. Nucleotide corresponds to the coding of the gene regardless of the translation product. Protein ref. is the PDB reference to access to all the information about that actin. Amino acid (green-magenta) and nucleotide (blue-orange) identities are at the end of the table. Positive is similar to identity but instead of considering all mutations in the protein, it considers only the non-conservative substitutions, meaning the changes in amino acids that have different chemical properties.

We synthesized the actin nucleotide sequences and sub-cloned them in a plasmid created specifically for rapid and robust actin gene replacement under endogenous promoter control in *S. cerevisiae* (Fig. S1D). With this strategy, we created a library of yeast strains, from which we systematically studied the effect of deleting the actin intron in haploid cells, changing the nucleotide sequence without modifying the final actin protein in haploid cells, switching actin protein variants in haploid cells, and expressing copolymers of actin in diploid cells (Fig. 1 D).

Previous studies have demonstrated that the yeast actin intron is not essential for actin gene transcription and for normal cell growth (Ng *et al*, 1985). Indeed, our analysis found that an *act1* gene construct without the intron in *S. cerevisiae* S288C (ScNI) does not affect cell growth (Fig. S2 A and B) nor actin expression (Fig. S2 C and D). Fixation and phalloidin-labeling of the actin cytoskeleton reveals that the two main structures of actin filaments in yeast, actin patches and actin cables, are well-organized in yeast strains expressing actin in the absence of the intron and indistinguishable from wild type cells (Sc) (Fig S2, E-G). Therefore, all experiments presented in the following sections of this study were conducted on actins expressed in the absence of an intron.

### Cell fitness tolerates reduced wild-type actin expression above a threshold

We were concerned that small changes to the actin nucleotide sequence might have consequences on actin expression levels and cell viability (Hoekema *et al*, 1987; Zhou *et al*, 2016). In mammals, for instance, nucleotide sequence was shown to differentiate beta and gamma actin functions (Vedula *et al*, 2017). Therefore, we expressed wild-type actin from a range of different nucleotide sequences. We used coding sequences from other organisms, which we modified minimally so that the final product remained *S. cerevisiae*’s actin at the protein level (Table 1) (Fig. S2 H). Western blot analysis showed that silent mutations affect wild-type actin’s expression level to various extents (Fig. 2, A and B), with a clear correlation between actin expression and the level of conservation of the nucleotide sequence (Fig. S2 I). These data also revealed that a sizeable drop of actin expression (for example, Act_Sc[Sp], derived from S. pombe’s actin gene, is expressed at 35% of normal level), has little or no effect on cell viability (Fig. 2, C-E and S2 J) nor on the organization (Fig. 2, F-H) or polarity (Fig. 2 I) of the actin cytoskeleton. However, a more drastic drop of actin expression (for example, Act_Sc[At], derived from *A. thaliana*’s ACT8 gene, is expressed at 24% of normal level), affects visibly cell viability (Fig. 2, C-E and S2 J), the organization (Fig. 2, F-H) and the polarization (Fig. 2I) of the actin cytoskeleton. Expressing actin from a gene derived from the nucleotide sequence of *H. sapiens* ActB, whose nucleotide sequence is even less conserved, is lethal for cells. From these observations, we concluded that the level of expression levels of actin orthologs should be controlled carefully in this study. Nevertheless, these results also indicated that variations in actin expression down to a ~ 35% threshold generally have negligible effect on cell behavior.

**Figure 2.**
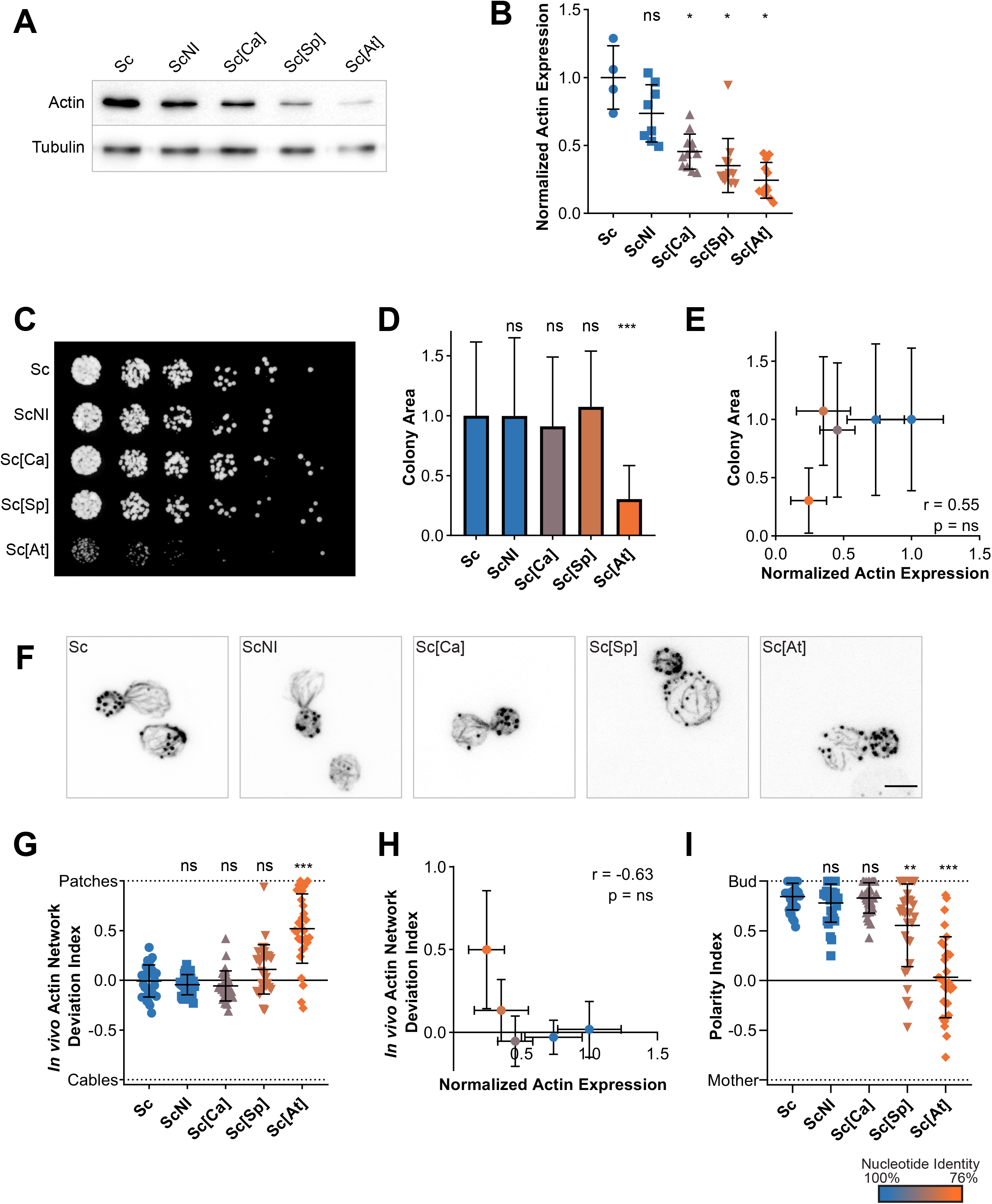
Effects of silent mutations on actin expression levels, cell viability and cytoskeletal organization. **(A)** Actin expression levels shown by western blotting for strains expressing *S. cerevisiae*’s actin protein from various coding sequences, with tubulin (Tub1p) as a loading control. **(B)** Quantification of actin expression levels, showing a decrease when more silent mutations are present. **(C)** 3-fold serial dilutions of yeast strains cultures, grown at 25 °C for 2 days on a YPD plate. **(D)** Quantification of (C) by measurement of colony area. **(E)** Level of actin expression as a function of colony area does not show any clear correlation. Rather, there is an apparent level of actin expression (0.25 < expression < 0.35) below which growth rates drastically reduce. **(F)** Phalloidin staining depicting F-actin organization. Images are maximum intensity projections of 3D stacks. Scalebar: 3 μm. **(G)** *In vivo* actin network deviation indexes, defined to evaluate the patch-cable balance compared to *S. cerevisiae* haploid cells (value is 0 in *S. cerevisiae*’s cells, 1 when cells contain only actin patches and −1 when cells contain only cables). **(H)** *In vivo* actin network deviation indexes as a function of actin expression levels does not show any clear correlation. Rather, we observe a threshold of actin expression levels (0.25 < expression < 0.35) below which actin cytoskeleton organization is affected. **(I)** Polarity indexes, defined to assess whether cell polarity is normal or affected (value is 1 when all patches of medium to large budded cells are present in the bud, and −1 refers when all patches are in the mother cell. Color code: nucleotide sequences percentage identities compared to *S. cerevisiae* actin gene, ranging from 100% (blue) to 75% (orange). Abbreviations: Sc - wild-type *S. cerevisiae* cells, ScNI - *S. cerevisiae* cells where the actin gene has been replaced with the wild-type gene but without the intron, Sc[X] – *S. cerevisiae* cells where the actin gene has been replaced with a gene carrying silent mutations based on the sequences from species X (for the list of species and coding see Table 1 or Figure 1). Statistics: Brown-Forsythe and Welch ANOVA tests, with Dunnett’s T3 multiple comparisons tests. P value style: GP <0.05, ** <0.01, *** <0.001. Error bars indicate standard deviations. Correlation coefficients r correspond to Pearson correlation coefficients.

### Actin amino acid sequence variations affect cell fitness and imbalance the linear-to-branched actin network ratio

We next focused our attention on the consequences of expressing heterologous actin orthologs in yeast cells. Actin genes were designed based on *S. cerevisiae*’s Act1 sequence by making point mutations using yeast codon usage. Overall, all coding sequences used in this section are more than 90% identical to that of *S. cerevisiae*, which according to the previous section, lowers the risk that actin expression is reduced excessively. Nevertheless, we verified the expression level of each heterologous actin ortholog in haploid strains by western blotting. The expression level of each actin varied, and appeared not to be correlated with the evolutionary relationship (Fig. 3, A-B). For example, Act_N1 was only expressed at 39% despite having a 98.4% identity to wild-type actin and showed normal viability and cytoskeletal organization (Fig. 3, C-G).

**Figure 3.**
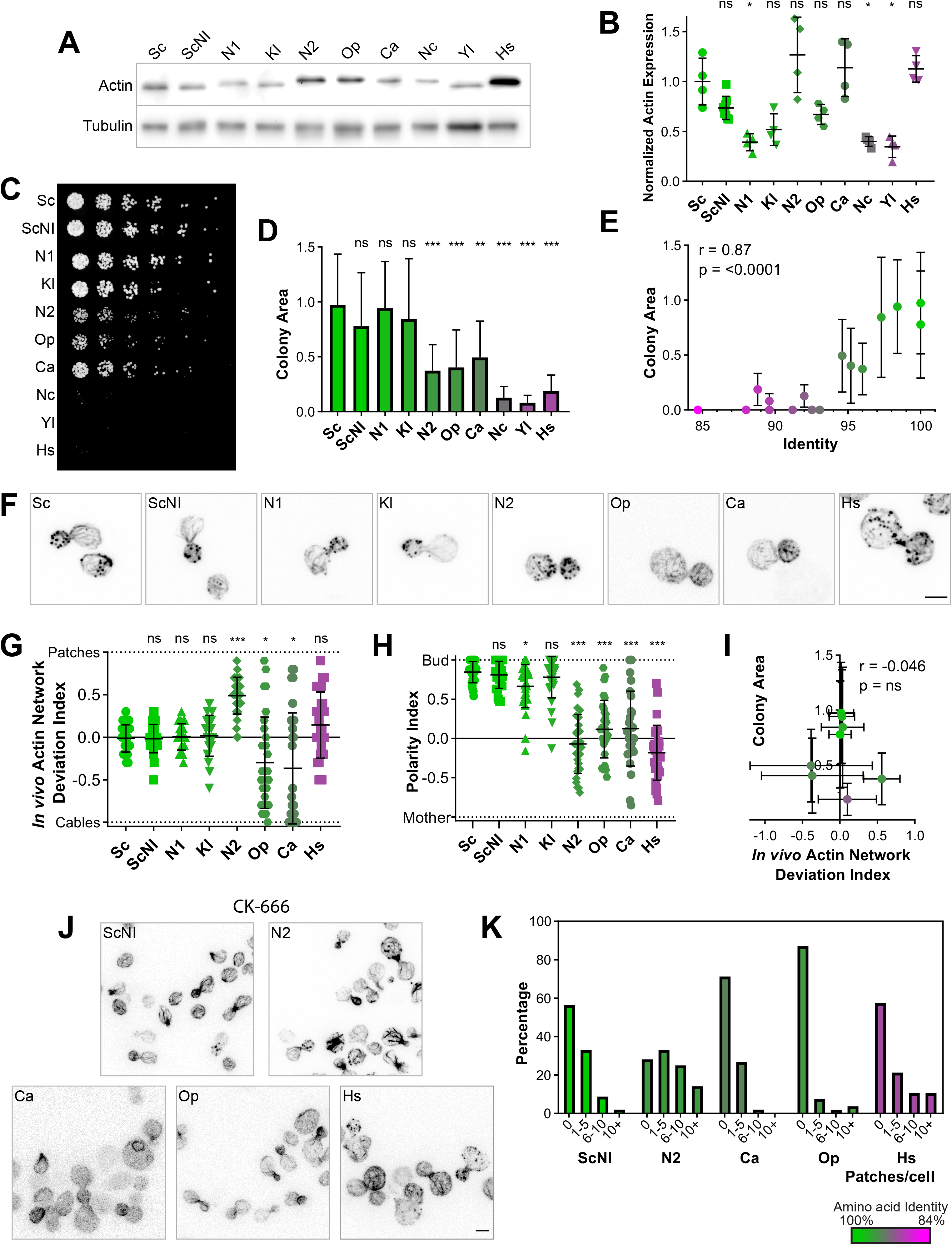
Effects on cell viability and cytoskeletal organization of swapping actin for different variants. **(A)** Actin expression levels shown by western blotting for strains expressing *S. cerevisiae*’s actin or other actins, with tubulin (Tub1p) as a loading control. **(B)** Quantification of actin expression levels showing varying levels of expression that do not correlate with evolutionary relationship. **(C)** 3-fold serial dilutions of different yeast strain cultures grown at 25°C for 2 days on a YPD plate. **(D)** Quantification of (C) by measurement of the colony area. **(E)** Colony area as a function of percentage identity of the actin variant, showing clear correlation. **(F)** Phalloidin staining of F-actin organization. Images are maximum intensity projections of 3D stacks. Scalebar: 3 μm **(G)** *In vivo* actin network deviation indexes. **(H)** Polarity indexes. **(I)** Colony area as a function of the *in vivo* actin network deviation index. **(J)** Effect of CK-666 (150 μM) on the organization of the actin cytoskeleton. Cells were stained with phalloidin after 30 min incubation with CK-666. Images are maximum intensity projections of 3D stacks. **(K)** Quantification of actin patch resistance to CK-666 treatment. Bar graphs represent the percentage of cells with a given number of visible actin patches after CK-666 treatment. Color code: nucleotide sequences percentage identities compared to *S. cerevisiae* actin, ranging from 100% (blue) to 84% (orange). Abbreviations: Sc - wild-type *S. cerevisiae* cells, ScNI - *S. cerevisiae* cells where the actin gene has been replaced with the wild-type gene but without the intron, the other abbreviations correspond to cells expressing actins from other species (for the list of species see Table 1 or Figure 1). Statistics: Brown-Forsythe and Welch ANOVA tests, with Dunnett’s T3 multiple comparisons tests. P value style: GP * <0.05, ** <0.01, *** <0.001. Error bars indicate standard deviations. Correlation coefficients r correspond to Pearson correlation coefficients.

While yeast strains expressing heterologous actin orthologs similar to *S. cerevisiae* wild-type actin (identity > 97%) grew well (Fig. 3, C-E), had normal cytoskeletal organization (Fig. 3, F-G) and were polarized normally (Fig. 3H), yeast strains expressing more distant actins (identity < 97%) showed moderate to severe defects (Fig. 3, C-H). The strength of the growth phenotypes correlated with the degree of conservation of the actin orthologs (Fig. 3E). Interestingly, consequences on the organization of the actin cytoskeleton was not the same for all mutants. While cells expressing Act_N2 assembled, on average, an abnormally high number of actin patches, strains expressing Act_Op or Act_Ca assembled, on the contrary, a higher number of actin cables and few actin patches (Fig. 3, F-G, I). Considering that actin networks do not assemble independently in cells, but that homeostatic actin networks share a limited monomeric actin pool (Burke *et al*, 2014), the previous observation suggests an imbalanced assembly between branched- and linear-actin structures from the use of different actin variants. We hypothesized that the cellular machinery cannot use Act_N2 efficiently to assemble actin cables, and cannot use Act_Op or Act_Ca to assemble actin patches, thus leading to an overproduction of patches in Act_N2 cells and an overproduction of cables in Act_Op and Act_Ca cells. It is also possible that patch or cable assembly is boosted by the use of a particular actin ortholog, although this hypothesis seems less likely, since it is generally easier to disrupt a function than make it more efficient.

Following this hypothesis, we tested whether strains over-assembling actin patches would show acute resistance to Arp2/3 perturbations. This was the case, as the strain expressing Act_N2 showed persistence of actin patches on treatment with the small molecule inhibitor of the Arp2/3 complex CK-666 (Fig. 3J,K). Conversely, Act_Op and Act_Ca strains were more sensitive to CK-666. These results indicate that strains with increased branched network are buffered against Arp2/3 perturbations.

### A biomimetic assay recapitulates actin ortholog preference for branched- or linear-network assembly

We then aimed to understand the molecular principles that allow different actin ortholog to be assembled specifically to certain actin networks, and hypothesized that heterologous actin orthologs may bind defectively to certain ABPs of *S. cerevisiae*. Because actin assembly into patches and cables involves a large number of proteins in cells, we adopted a reductionist approach based on a reconstituted assay. We considered that the subset of ABPs that are most essential for actin patch or cable assembly *in vivo*. Beyond formins and the Arp2/3 complex, these proteins include: profilin, a small globular protein that favors formin assembly, capping protein, a heterodimer that binds to barbed ends, ADF/cofilin, a small protein that promotes the disassembly of actin filaments, and tropomyosin, a helical coiled-coil protein that binds and stabilizes linear-actin filaments nucleated by formins (Moseley & Goode, 2006; Pollard, 2016).

In addition to wild-type actin, we purified Act_N2 and Act_Ca from cultures of the corresponding yeast strains. We reconstituted *in vitro*, in a common experimental environment, branched- and linear-actin network assembly, respectively from formin- and WASp-coated beads (Antkowiak *et al*, 2019). First, we assessed the capabilities of the three actin orthologs to assemble into such networks. Act_Ca assembled only into linear-actin networks (Fig. 4A), providing explanation for the inability of Act_Ca cells to assemble actin patches. However, Act_N2 assembled both into branched- and linear-actin networks similarly to the control condition. We therefore hypothesized that an ABP, involved in the stabilization or disassembly of one of those actin networks, may bind abnormally. We labeled ADF/cofilin, which is known to promote branched-network disassembly by inducing Arp2/3 debranching while stabilizing linear-networks (Michelot *et al*, 2007; Chan *et al*, 2009). ADF/cofilin bound to linear-actin networks with higher affinity than to the branched-actin networks (Fig. 4B), as previously reported (Gressin *et al*, 2015). However, ADF/cofilin bound similarly to both actin variants, albeit with reduced affinity compared to wild-type actin (Fig. 4B). We next labeled tropomyosin, which inhibits branched-network assembly and promotes linear-network stabilization (Blanchoin *et al*, 2001; Bernstein & Bamburg, 1982; DesMarais *et al*, 2002; Antkowiak *et al*, 2019). Tropomyosin bound with higher efficiency to linear-actin networks, as expected (Fig. 4C). Its binding to Act1p and Act_Ca was similar; however, tropomyosin was completely absent from actin networks assembled from Act_N2 (Fig. 4C). This inability to bind to Act_N2 provides a likely explanation why actin patch assembly is favored in Act_N2 cells.

**Figure 4.**
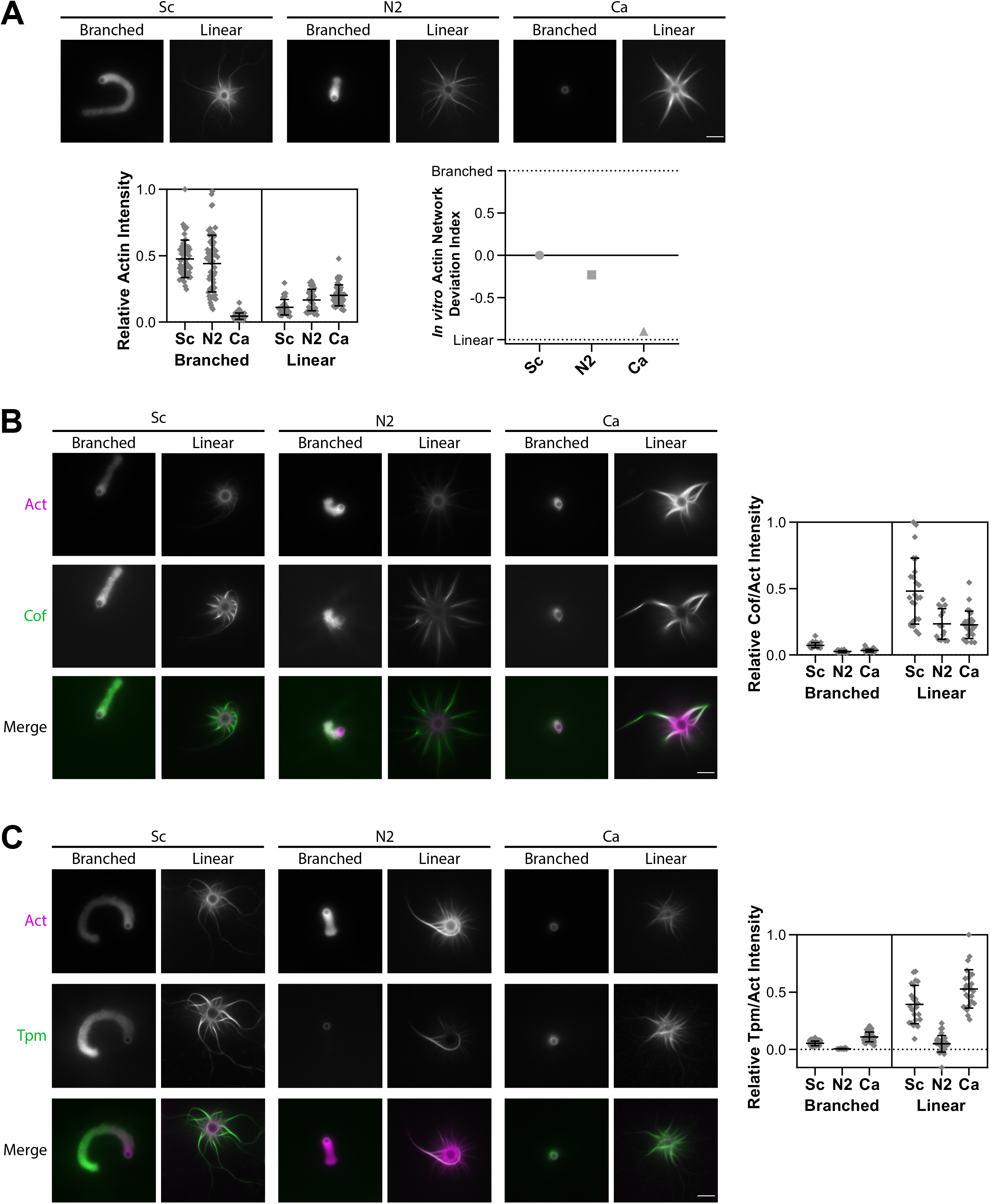
*In vitro* reconstitution of branched- and linear-actin networks assembly from purified actins. Standard conditions include Las17- (branched) and Bni1- (linear) coated beads, 8 μM F-actin (1% Alexa-568-labeled), 15 μM profilin, 1 μM capping protein, 500 nM Arp2/3 complex and 600 nM ADF/cofilin. Snapshots of representative actin networks were taken after 30 min. Scalebars: 6 μm. **(A)** (Top) Snapshots of actin networks assembled from three different actins sources: Act1, Act_N2 and Act_Ca. (Bottom left) Quantification of actin fluorescence on beads. (Bottom right) *In vitro* actin network deviation indexes. **(B)** (Left) Snapshots of representative actin networks assembled in the presence of 600 nM Alexa-488-labeled ADF/cofilin (replacement of unlabeled ADF/cofilin). **(C)** Snapshots of representative actin networks assembled in the presence 1 μM Alexa 488-tropomyosin. For all microscopy images, contrasts were adapted for images of branched- and linear-actin networks separately as their brightness is different. Please refer to quantifications on the left to compare levels of intensity. Abbreviations: Sc – purified *S. cerevisiae* actin, N2 – purified Node 2 actin, Ca – purified *C. albicans* actin (for more details see Table 1, Figure 1 and Figure S1 B-C).

### Structural analysis provides plausible explanation of defective interactions

We searched for a structural understanding of why Act_N2 and Act_Ca do not interact properly with specific ABPs of *S. cerevisiae*. Based on the structural information available of the interactions of actin with its binding partners, we identified actin residues that are within 5 Å of at the actin-actin interface in a filament, or at the interface between G- or F-actin and the ABPs used in our biomimetic assay (Winn *et al*, 2011), with the exception of the Arp2/3 mother filament which were within 10 Å since the coordinates were not released when this study was performed (Fäßler *et al*, 2020) (Fig. 5A).

**Figure 5.**
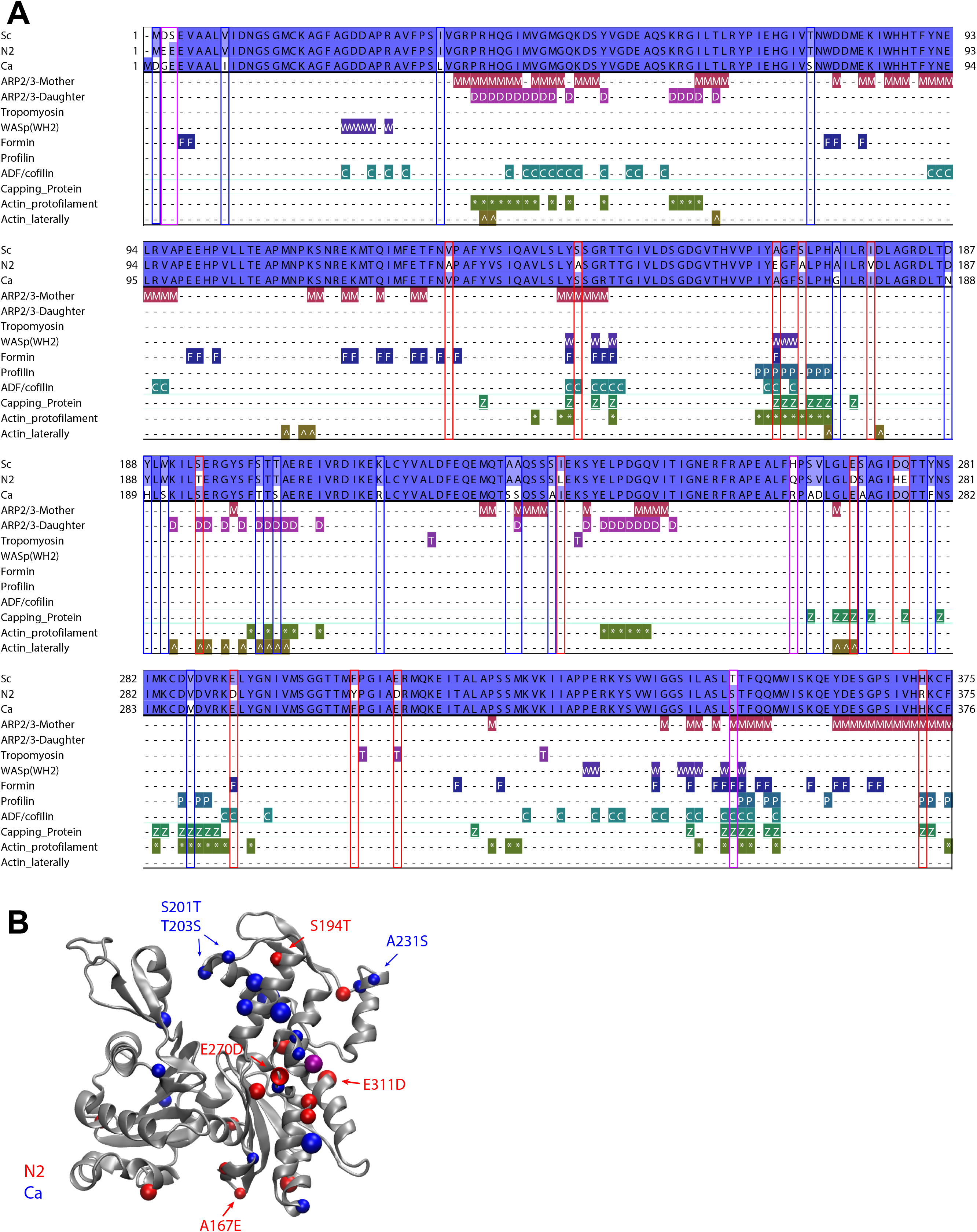
ABPs interfaces with actin. **(A)** Sequence alignment of three actins (Act_Sc, Act_N2 and Act_Ca; deep blue indicates conserved residues, light blue and white indicates non-conserved), indicating contacts between proteins used in the biomimetic assay (with Arp2/3 complex at the mother filament interface (M), with Arp2/3 complex at the daughter filaments interface (D), with tropomyosin (T), with WASP’s WH2 (W), with formin (F), with profilin (P), with ADF/cofilin (C), with capping protein (Z), at the protofilament interface (*) and laterally (^). **(B)** Schematic representation of actin 3D structure (1YAG (Vorobiev *et al*, 2003)). Color dots correspond to positions where Act_Sc has different residues compared to Act_N2 (red) and Act_Ca (blue). Purple dots correspond to positions where both Act_N2 and Act_Ca have different residues compared to Act_Sc. Abbreviations: Sc –*S. cerevisiae* actin, N2 – Node 2 actin, Ca – *C. albicans* actin (for more details see Table 1, Figure 1 and Figure S1 B-C).

At protomer:protomer interfaces, wild-type actin differed by one residue (Val287Met) and two residues (Ala167Glu and Ser170Ala) relative to Act_Ca and Act_N2, respectively (Fig. 5B). In particular, the Ala167Glu substitution has been shown to effect actin filament stiffness (Hocky *et al*, 2016; Kang *et al*, 2012; Scipion *et al*, 2018). Furthermore, four differences were observed in inter-strand contacts relative to Sc (Ser194Thr and Glu270Asp for Act_N2) and (Ser201Thr and Thr203Ser for Act_Ca) (Fig. 5B). Together, these substitutions may subtly alter the relative filament plasticity, which in turn may have an influence on the association or activity of filament binding and filament nucleating proteins (McCullough *et al*, 2011; von der Ecken *et al*, 2015). In addition, we identified 15 non-conserved residues of Act_N2 or Act_Ca that are surface exposed on the actin protomer structures and contact a binding partner (Table 2). Tropomyosin is likely to be particularly susceptible to small changes in the actin filaments, since it loosely associates with the actin filament surface via shape and charge complementarity (Popp & Robinson, 2012; von der Ecken *et al*, 2016). Particularly, Act_N2 filament Asp311 potentially places the negative charge at ~1.5 Å closer to the actin, relative to the Sc and Ca filaments (glutamic acid), which may be inappropriate for tropomyosin binding. Act_N2 has substitutions in interfaces with all the proteins used in the *in vitro* assays, including Arp2/3 and formin interfaces, which could have impaired the activities of these filament nucleating complexes. Act_Ca has fewer substitutions in the actin regulating proteins, with the notable exception of Arp2/3. In particular, substitutions in the actin interfaces with Arp2/3 subunits in the daughter filament may indicate that the nucleation process of the daughter filament is impaired for Act_Ca with *S. cerevisiae*’s Arp2/3.

**Table 2.** ABPs interfaces with actin, identified in CONTACT (Winn *et al*, 2011). Detail of the mutations in Act_N2 (red) and Act_Ca (blue) compared to Act_Sc (black) and the interaction of each position with ABPs. Abbreviations: Sc –*S. cerevisiae* actin, N2 – Node 2 actin, Ca – *C. albicans* actin (for more details see Table 1, Figure 1 and Figure S1 B-C).

| *Saccharomyces cerevisiae* | Node 2 | *Candida albicans* | Actin Proto | Actin Lateral | Capping Protein | Profilin | Cofilin | Arp2/3 Daughter | Arp2/3 Mother | Formin | WH2 | Tpm |
| --- | --- | --- | --- | --- | --- | --- | --- | --- | --- | --- | --- | --- |
| Ser144 | Ala144 | Ser145 |  |  |  |  | N2 |  | N2 |  |  |  |
| Ala167 | Glu167 | Ala168 | N2 |  | N2 | N2 | N2 |  |  | N2 | N2 |  |
| Ser170 | Ala170 | Ser170 | N2 |  |  |  |  |  |  |  |  |  |
| Ser194 | Thr194 | Ser195 |  | N2 |  |  |  | N2 |  |  |  |  |
| Ser201 | Ser201 | Thr202 |  | Ca |  |  |  | Ca |  |  |  |  |
| Thr203 | Thr203 | Ser204 |  | Ca |  |  |  | Ca |  |  |  |  |
| Ala232 | Ala232 | Ser234 |  |  |  |  |  | Ca | Ca |  |  |  |
| Ser265 | Ser265 | Ala266 |  |  | Ca |  |  |  |  |  |  |  |
| Glu270 | Asp270 | Glu271 |  | N2 | N2 |  |  |  |  |  |  |  |
| Gln276 | Glu276 | Gln277 |  |  | N2 |  |  |  |  |  |  |  |
| Val287 | Val287 | Met288 | Ca |  | Ca |  |  |  |  |  |  |  |
| Glu292 | Asp292 | Glu293 |  |  |  |  | N2 |  |  | N2 |  |  |
| Glu311 | Asp311 | Glu312 |  |  |  |  |  |  |  |  |  | N2 |
| Thr351 | Ser351 | Ser315 |  |  | N2/Ca |  | N2/Ca |  | N2/Ca | N2/Ca |  |  |
| His372 | Arg372 | His373 |  |  | N2 | N2 |  |  | N2 |  |  |  |

### Dual expression of a patch and a cable-favoring actin rescues cell viability and cytoskeletal organization

The identification of heterologous actin orthologs favoring the specific assembly of actin patches or cables suggested that actin functions could be separated from the use of two carefully selected actin variants (Fig. 6A). To test this possibility, we switched to a diploid yeast cell background. We verified first that both Act_N2/Act_N2 and Act_Ca/Act_Ca cells display similar phenotypes to their haploid equivalents, with slow growth and unbalanced actin patch and cable assembly (Fig. 6B-C). We then crossed strains in order to express a copy of each actin variant in the same cell. Strikingly, cell growth (Fig. 6B-C), actin cytoskeleton organization (Fig. 6D-E) and cell polarity (Fig. 6F) were rescued in diploid cells expressing both Act_N2 and Act_Ca. Verification that each of the actin structures was enriched by each of the variants is difficult to do in the absence of specific antibodies; nevertheless, our results indicate that defective actin functions in cells carrying a single actin variant were carried out more normally when the other actin variant was simultaneously expressed.

**Figure 6.**
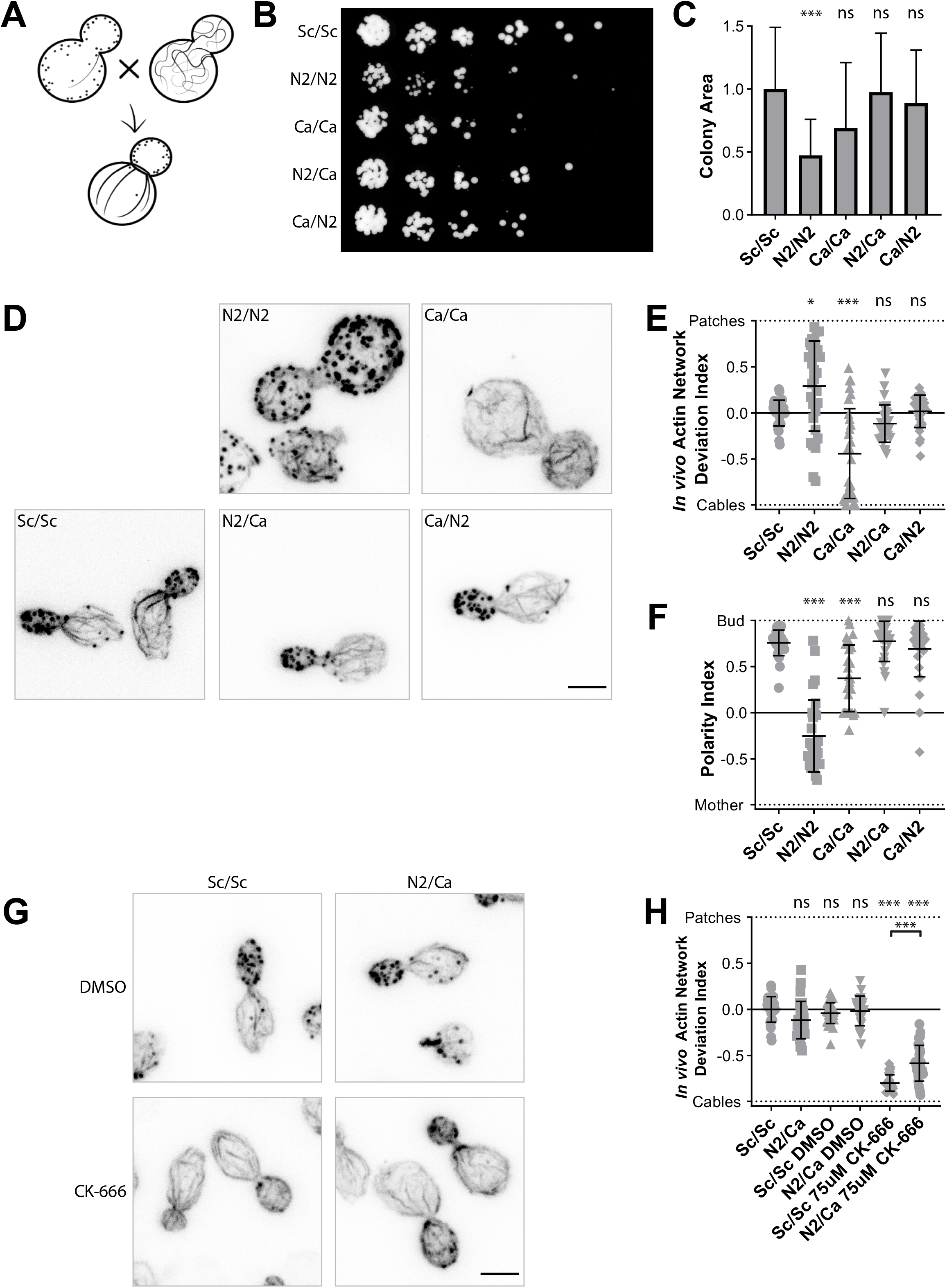
Effect of a dual expression of actins on cell viability and cytoskeletal organization. **(A)** Schematic of the experiment performed in a diploid yeast background. **(B)** 3-fold serial dilutions of diploid yeast strains cultures, grown at 25°C for 2 days on a YPD plate. Act_N2/Act_Ca and Act_Ca/Act_N2 express the same actins but markers used for selection are exchanged. **(C)** Quantification of (B) by measurement of colony area. **(D)** Phalloidin staining depicting F-actin organization. Images are maximum intensity projections of 3D stacks. **(E)** *In vivo* actin network deviation indexes. **(F)** Polarity Indexes. **(G)** Effect of CK-666 (75 μM) on the organization of the actin cytoskeleton. Cells were stained with phalloidin after 30 min incubation with CK-666. Images are maximum intensity projections of 3D stacks. **(H)** *In vivo* actin network deviation indexes of cells treated with DMSO or CK-666. Scalebars: 3 μm. Abbreviations: Sc/Sc – wild-type diploid *S. cerevisiae* cells, N2/N2 – diploid *S. cerevisiae* cells expressing only N2 actin, Ca/Ca – diploid *S. cerevisiae* cells expressing only *C. albicans* actin, N2/Ca and Ca/N2 – diploid *S. cerevisiae* cells expressing N2 actin and *C. albicans* actin at the same time (for more details see Table 1, Figure 1 and Figure S1 B-C). Statistics: Brown-Forsythe and Welch ANOVA tests, with Dunnett’s T3 multiple comparisons tests. P value style: GP * <0.05, ** <0.01, *** <0.001. Error bars indicate standard deviations.

### F-actin network homeostasis is affected in a two-actin system

Generation of yeast strains with partially separated actin functions enabled us to question some differences between species sharing a single actin for multiple cellular functions, and species using different actin variants. We were especially curious to know what the physiological consequences would be on actin network homeostasis for wild type diploid cells and Act_N2/Act_Ca cells, which share the same ratio of branched and linear network but possess different actin variants. As expected, addition of CK-666 in wild-type cells resulted in the disappearance of actin patches and an increase of actin cables (Fig. 6G-H). On the contrary, addition of CK-666 to Act_N2/Act_Ca cells had a weaker effect, as a large number of actin patches could still be observed (Fig. 6G-H). Together, these results show that while F-actin network homeostasis is preserved in a yeast strain using a single actin ortholog, actin re-distribution from one network to another is less effective in the context of a yeast strain expressing two different actin variants.

## Discussion

### Identification of actin variants that favor branched- or linear-actin networks assembly

He we investigated, at the cellular and at the molecular level, the consequences of perturbing a simple system which uses a single actin ortholog and a limited number of regulatory proteins to assemble an organized actin cytoskeleton. We demonstrated that small variations in the actin sequence are sufficient to induce a global reorganization of the actin cytoskeleton. This finding highlights the fact that, despite the remarkably high sequence conservation of actin orthologs across species, there are sufficient differences in sequence for cells to segregate multiple actin variants into diverse actin networks.

Generally, we found that mutant cells expressing a heterologous actin assemble an abnormal distribution of actin patches and cables. This result is coherent with the literature, which shows particularly in yeast that homeostatic actin networks compete for a limited pool of actin monomers (Burke *et al*, 2014) (Fig. 7 top and bottom left). In this context, it is rational to postulate that the inability of an actin variant to assemble efficiently in a given actin network, leads to an expansion of the other actin networks, provided that those can use this actin normally (Fig. 7 top middle).

**Figure 7.**
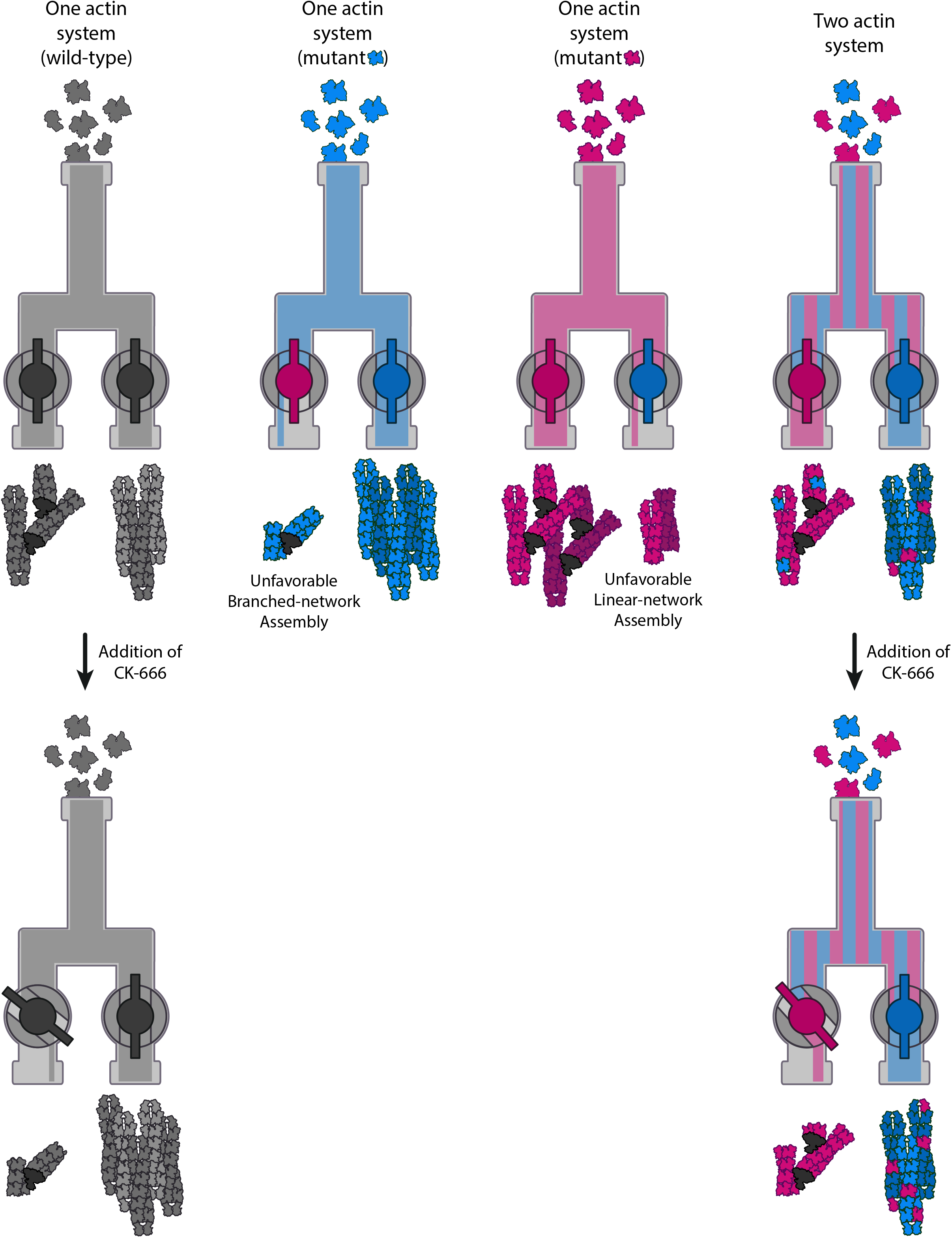
Schematic model of the differences between a cell expressing one or two actins to perform two cellular functions. **(Top)** A model of the molecular mechanisms by which two actin isoforms may segregate to different actin networks. On the left, a system carrying wild-type actin is able to generate both the branched- and linear-networks. On the two central panels, defective interactions of an actin isoform with one or several ABPs, affect branched- or linear-network assembly. On the right, combining these two actin variants in one cell should trigger a natural segregation of actins and rescues the wild type actin organization. **(Bottom)** Effect of perturbing an actin assembly pathway for cells using one or two actin variants. On the left, when one actin is shared for two actin functions, the inhibition of one actin assembly pathway (for example branched-networks with CK-666) leads to a reinforcement of the other actin assembly pathway. On the right, when two actin variants are used for two different actin functions, this effect is limited as both actin networks assemble more independently. In other words, having a system with two actin variants can buffer against the addition of the drug.

The possibility to rescue yeast cell viability with the simultaneous expression of patch-favoring and cable-favoring actin variants, reinforces the possibility that actin isoforms compensate for the other actin’s lack of efficiency to form a certain structure (Fig. 7 top right). The lack of specific probes that can differentiate between these actin variants prevented us from localizing them in cells and from verifying the extent of segregation. We expect the integration of an actin variant within a particular network to be dependent on its innate ability to assemble in such a network, and also to be affected by relative efficiencies of other co-expressed actin variants to integrate within branched- and linear- actin networks. Such a hypothesis is purely speculative and should be formally tested in the future. Nevertheless, our observations strongly suggest that we were successful in performing a partial separation of function, and in transforming yeast from being an organism that uses a single actin into an organism using two actin variants to perform several actin-based functions.

### Molecular subtleties guide actins to appropriate networks

This study was originally motivated to provide a systematic description of the molecular principles by which different actin isoforms could become spatially segregated into different networks. Experiments performed here show that a biomimetic system, using a reduced set of essential proteins for patch and cable assembly, is sufficient to provide basic molecular explanation of differences observed in cells. While one actin (Act_Ca) seems inefficient in nucleating or assembling into branched-actin networks, the other actin (Act_N2) seems to assemble in both types of actin networks. However, Act_N2 is defective in binding to tropomyosin, which is an essential component for cable stability in cells, as it protects them from the action of disassembly factors such as ADF/cofilin. This result highlights that the segregation of actin isoforms can be influenced after filament nucleation. Although actin nucleators tend to be in the spotlight, it must be stressed that most ABPs have different effects on branched- and linear-actin networks and their influence needs to be taken into account (Rotty *et al*, 2015; Suarez & Kovar, 2016; Antkowiak *et al*, 2019). Proteins like tropomyosin and ADF/cofilin stabilize linear networks of actin filaments, while enhancing disassembly of branched networks. In this context, any actin variant with defective binding to ADF/cofilin or tropomyosin will naturally be more present within branched networks, while absent from linear ones.

Overall, the principles outlined above should be valid regardless of the mechanism by which variation is brought to the specific actin, whether it is through changes in the peptide sequence or through post-translational modifications. Also, from an evolutionary perspective, the proposed mechanism appears to be efficient in allowing the emergence of new actin isoforms associated with discrete actin functions. Our model implies that a simple actin gene duplication, followed by minimal mutation in one actin copy, which impairs an essential interaction with an ABP, could be sufficient to trigger a global reorganization of the actin cytoskeleton, whereby each actin network becomes enriched in one actin isoform or the other.

We now have precise structural information on how actin interacts with many ABPs (Pollard, 2016; Tanaka *et al*, 2018; Fedorov *et al*, 1997; Shaaban *et al*, 2020; Baek *et al*, 2008; Eads *et al*, 1998; Urnavicius *et al*, 2015; Otomo *et al*, 2005; Thompson *et al*, 2013; von der Ecken *et al*, 2015). Careful analysis of actin-actin and actin-ABPs interactions can be a powerful tool to predict which ABPs affect the roles of specific actin isoforms in discrete actin networks. For example, such analysis indicates that most Act_Ca substitutions affect its interface with the Arp2/3 complex, providing potential explanation for defective assembly into branched-actin networks. In parallel, our knowledge of the molecular principles involved in the assembly of the different actin networks of the cell allows us to anticipate the consequences of varying the affinity between actin and an ABP.

### Consequences of multiple functions deriving from a single or multiple actin isoforms

Finally, the generation of a yeast strain carrying two different actin variants allowed us to question the main differences between systems using the same actin and systems using several actin isoforms to perform various actin-based functions. We showed that addition of CK-666 in strains expressing both actins Act_Ca and Act_N2 did not lead to similar cytoskeletal reorganization as in wild-type strains (Fig. 7 bottom). While cells expressing wild-type actin can easily shift actin use from patches to cables, the mechanism was less efficient for a two-actin system, indicating a perturbed homeostasis of actin networks. This observation brings additional evidence that assembly of both actin networks is more independent in a two-actin system. Therefore, it is possible that an important difference highlighted here is that organisms using a single actin for multiple actin functions have the possibility for global reorganization of the actin cytoskeleton, where increased assembly of a specific actin network occurs at the expense of others. Conversely, for organisms using multiple actin isoforms, the various actin assembly pathways may be modulated separately, allowing for more autonomous actin networks and functions.

## Material and methods

### Reconstruction of ancestral protein sequences and selection of actins

Actin amino acid sequences from 122 different species, selected from different branches of the eukaryotic tree of life to cover a wide range of variations, were collected from anotated and reviewed UniProtKB/Swiss-Prot entries. For species encoding more than one actin, the cytoplasmic actin with the most similar sequence to *S. cerevisiae* actin was selected. The resulting 122 selected sequences from different species were aligned using the Multiple Sequence Alignment program Clustal Omega (Madeira *et al*, 2019) in Pearson/FASTA format. The phylogenetic tree of the 122 species was created based on the NCBI taxonomy using the phylogenetic tree generator phyloT (https://phylot.biobyte.de/). Ancestral actin sequence reconstruction was performed from multiple sequence alignment and phylogenetic tree inputs using FastML (Ashkenazy *et al*, 2012). 99% of the amino acids in the ancestral sequences are predicted with an accuracy >95% and uncertain residues correspond to conservative substitutions (Grantham score <100, (Grantham, 1974)).

### Generation of plasmids for efficient and rapid actin gene replacement

We generated two plasmid backbones, which carried in succession a sequence upstream of the yeast actin promoter (−804 to −467 from act1 gene) as a first site for homologous recombination, a first selection marker (URA3 or HIS3), the yeast actin promoter (−473 to 0), the *act1* coding sequence, a short sequence downstream of the actin gene (+1437 to +1703), a second selection marker (LEU2 or KanMX3) and lastly a sequence downstream of the yeast actin gene as a second site for homologous recombination (+1543 to +2071). The advantage of having two different selection markers within the same plasmid is to easily select correct insertions of DNA fragments from partial insertions which are more frequent when targeting an essential gene like actin. These plasmids also contain four unique restriction sites: PacI and XbaI, on each side of the actin gene, allowed to sub-clone easily new actin coding sequences in the plasmid; Bsu36I and AatII before and after the two sites for homologous recombination, allowed to obtain linear DNA fragments for yeast transformation.

The new actin genes used in this study were obtained commercially from whole gene synthesis techniques (Synbio Technologies). For analysis of *S. cerevisiae*’s actin expression effects from various nucleotide sequences, we selected multiple actin nucleotide sequences from the specified species and we point mutated the corresponding codons so that the translation product is *S. cerevisiae*’s actin. For analysis of exogenous actin expression effects, we manually changed the coding sequence of *S. cerevisiae*’s actin gene (*act1*) by changing the specific codons that correspond to amino acid mutations respecting the budding yeast codon usage. All plasmids generated for this study are listed in Table S2 and the actin sequences are given in Fig. S1 C.

### Yeast strain generation

Actin gene replacement was performed in diploid cells. *S. cerevisiae* were transformed using the LiAc/SS carrier DNA/PEG method (Gietz & Schiestl, 2007) and grown on dual selection media. Correct insertion of DNA fragments were verified by PCR for all strains and sequenced. Strains were stored as heterozygous diploids and haploid mutant strains were isolated by tetrad analysis for study. Tables S3 and S4 list all the haploid and diploid yeast strains used in this study.

### Yeast growth assays on plates

For yeast growth assays, yeast cells were grown in YPD (2% bacto-peptone, 1% yeast extract, 2% dextrose) overnight at 25°C. Equal amounts cells were calculated from log phase growing cultures were serially diluted and spotted on YPD plates. Pictures of plates were taken after 2 days of growth at 25°C.

### Actin cytoskeleton organization in yeast

#### • Yeast cell phalloidin staining and imaging

Log phase cultures in YPD medium at 25°C were fixed with 4% formaldehyde for 2 h. For CK-666-sensitivity assays, cells were treated with the indicated concentration of CK-666 (Sigma-Aldrich SML0006) for 30 min before fixation. After fixation, cells were washed twice in PBS and stained overnight with 250 nM Phalloidin-Alexa568 (Invitrogen, ref. A12380) at 25°C. Samples were washed twice with PBS, re-suspended in PBS-70% glycerol and directly mounted for imaging. Cells were imaged using a Leica TCS SP8 X White Light Laser confocal microscope equipped with a HC PL APO CS2 100x/1.4NA Oil objective and a hybrid detector. Z-stack images were collected every 0.3 μm with Las X 3.5.5.19976 software.

#### • Data analysis for live imaging

Branched- and linear-actin network assembly in medium budded cells was assessed from the intensity of actin patches and cables, respectively. Total cell cable intensities were calculated from maximum intensity z-stack projections using Fiji v.1.53a. For total cell endocytic patch intensities, patches were identified using the TrackMate plugin of Fiji (Planade *et al*, 2019; Tinevez *et al*, 2017). Patch detection was corrected manually using the spot editing tool and the integrated intensity for all patches was calculated from the analysis table of TrackMate.

Fluorescence intensity of phalloidin labeling varied between strains expressing different actin variants. For this reason, the contrast of images showed in the figures was adapted from strain to strain so that both actin structures remained clearly visible. In addition, rather than reporting total intensities, we compared the relative assembly of branched- and linear- actin networks for each strain. This choice is also motivated by the fact that actin networks do not assemble independently but compete for a limited pool of monomeric actin (Burke *et al*, 2014). We calculated an *in vivo* actin network deviation index, defined as in (Antkowiak *et al*, 2019):

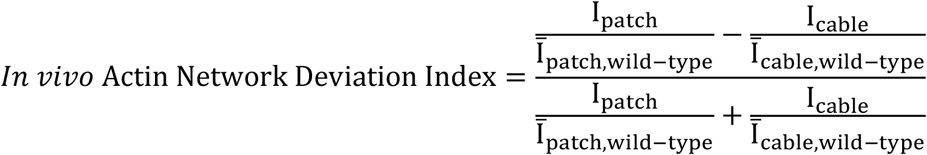

where I_patch_ (resp. I_cable_) is the total patch (respectively cable) fluorescence intensity of the cell of interest, and 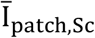 (resp. 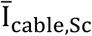) is the mean total intensity of actin patches (resp. cables) in wild type *S. cerevisiae*’s cells. This branched-to-linear actin network ratio was calculated for each cell, and compared to *in vivo* actin network deviation indexes of wild-type *S. cerevisiae*’s cells.

For cell polarity, number of visible patches in the bud (*Patches_bud_*) and in the mother cell (*Patches_mother_*) were taken into account to calculate a polarity index:

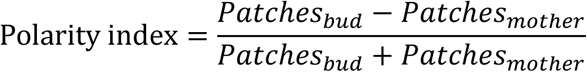

### Quantification of actin expression levels

A mouse anti-Actin C4 primary antibody (Fisher Scientific, ref. 08691002; 1:10,000 dilution) was selected to recognize actin. Its epitope is located around amino acids 50-70, which corresponds to a highly conserved region across all actins used in this study, with the exception of position 70 (a lysine in Homo sapiens beta actin; an arginine for all other actins). To test the antibody’s sensitivity to this amino acid variability, different amounts of purified rabbit muscle actin, which contains a lysine in position 70 and purified budding yeast actin, which contains an arginine in position 70, were loaded on a 12% gel. After protein migration, gels were transferred to a nitrocellulose membrane, incubated with mouse anti-Actin C4 primary antibody (Fisher Scientific, ref. 08691002; 1:10,000 dilution) overnight at 4°C, then incubated with a goat anti-mouse HRP antibody (Jackson ImmunoResearch, ref. 115-035-146; 1:10,000 dilution) for 1 h at 25°C and revealed with Western lightning plus ECL reagent (PerkinElmer, Inc., ref. NEL104001EA). After the Western blots were imaged, membranes were incubated with Ponceau S for 10 minutes and imaged again. Immunostaining signals were compared relative to Ponceau staining signals. Value of 12 measurements indicated on average a 1.48-fold stronger signal for rabbit muscle actin compared to *S. cerevisiae’s* actin. This value was used afterwards as a normalization factor when comparing the expression of Act_Hs and the expression of other actins in yeast.

Total protein samples from *S. cerevisiae* strains were prepared by trichloroacetic acid precipitation as described in (Reid & Schatz, 1982). 12% SDS-PAGE gels were loaded with 15 μg of total protein sample, and transferred to a nitrocellulose membrane after protein migration. Actin was recognized by anti-Actin C4 primary antibody (Fisher Scientific, ref. 08691002; 1:10,000 dilution). For our loading control, we selected a rabbit anti-alpha tubulin primary antibody (Abcam, ref. ab184970; 1:20,000 dilution). Western blots were incubated with goat anti-mouse HRP (Jackson ImmunoResearch, ref. 115-035-146; 1:10,000 dilution) and goat Anti-rabbit IgG H&L (HRP) (Abcam, ref. ab205718; 1:20,000 dilution) secondary antibodies. Western blots were revealed with Western Lightning Plus ECL reagent (PerkinElmer, Inc., ref. NEL104001EA), on a ChemiDoc MP imaging system (BioRad). We verified the linearity of results obtained with this method over a range of 3 μg to 30 μg of total extract loaded in the gels. Bands intensities were calculated using the Image Lab 6.0.1 software. Actin signals were relativized to the tubulin signals. The same control sample was loaded in all membranes and all values were normalized to this lane.

### Protein purification and labeling

#### Actins

Strains expressing Act1, Act_N2 and Act_Ca were used to purify the respective actins. Large-scale cultures were prepared at 25°C in YPD and harvested by centrifugation. Pellets were frozen in liquid nitrogen and ground in a steel blender (Waring, Winsted, CT, USA) (Michelot & Drubin, 2014). Actins were affinity-purified on a DNAse I- column (Goode, 2002). Yeast powder was resuspended in G1 buffer (10 mM Tris-HCl, 0.5 mM ATP, 0.2 mM DTT, 0.2 mM CaCl_2_) containing protease inhibitors (Protease Inhibitor Cocktail Set IV, Calbiochem, reference 539136), and centrifuged for 30 min at 160,000 x g at 4°C. The lysate was passed through a DNase I column. Bound actin was purified and eluted with G1 buffer supplemented with 50% formamide, and dyalised against G1 buffer with less calcium (10 mM Tris-HCl, 0.5 mM ATP, 0.2 mM DTT, 0.1 mM CaCl_2_) overnight. Rabbit muscle actin was purified from standard procedures (Spudich and Watt, 1971).

#### Actin labeling

G-actin from rabbit muscle was dialyzed against 25 mM Hepes pH 7.5, 50 mM KCl, 0.1 mM CaCl_2_, 0.2 mM ATP) at 4°C for 12 h. A 6-fold-excess of Alexa Fluor 568 succinimidyl ester dye was added and incubated overnight. F-actin was then centrifugated at 390,000 x g for 40 min, pellet was resuspended and dialyzed against G buffer for 2 h at 4°C. Labeled actin was centrifugated at 390,000 x g for 40 min to remove insoluble components and labeled actin was eventually loaded into a G25 column to remove unbound fluorophore.

#### Formin

*S. cerevisiae* cells (MATa, leu2, ura3-52, trp1, prb1-1122, pep4-3, pre1-451) were transformed with a plasmid designed for formin overexpression (Gst-Bni1(1215-Cter)-TEV-9xHis) under the control of a GAL1 promoter) (Antkowiak *et al*, 2019)). The expression was induced with 2% galactose for 12 h at 30°C. The resulting cultures were centrifuged and cells were frozen in liquid nitrogen and ground in a steel blender. For protein purification, 5 g of yeast powder was thawed on ice with 45 ml of HKI10 buffer (20 mM Hepes, pH 7.5, 200 mM KCl, 10 mM imidazole, pH 7.5), supplemented with 50 μl of Protease Inhibitor Cocktail Set IV and centrifugated at 160,000 x g for 30 min. The supernatant was collected and then incubated with 500-μl of Nickel-Sepharose 6 Fast Flow (GE Healthcare Life Sciences, Piscataway, NJ, USA) for 2 h at 4°C. Protein bound to Nickel-Sepharose beads was washed with HKI20 buffer (20 mM Hepes, pH 7.5, 200 mM KCl, 20 mM imidazole, pH 7.5) and cleaved from the beads by a 1 h incubation with TEV at room temperature. The protein was concentrated with an Amicon Ultra 4 ml device (Merck4Biosciences), dialyzed against HKG buffer (20 mM Hepes, pH 7.5, 200 mM KCl, 6% glycerol), flash frozen and stored −80°C.

#### Arp2/3 complex

*S. cerevisiae* Arp2/3 complex was purified from commercially purchased baker’s yeast (L’Hirondelle) based on a protocol modified from (Nolen & Pollard, 2008; Doolittle *et al*, 2013; Antkowiak *et al*, 2019). Yeast powder was prepared by flash freezing droplets of liquid yeast culture in liquid nitrogen and grinding them in a steel blender. 230 g of yeast powder was resuspended in a lysis buffer (20 mM Tris-HCl pH 7.5, 150 mM NaCl, 2 mM EDTA, 1 mM DTT) supplemented with Protease Inhibitor Cocktail Set IV. The mixture was centrifuged at 160,000 x g for 30 min and the supernatant was fractioned by a 50% ammonium sulfate cut. The insoluble fraction was dissolved, dialyzed in HKME buffer (25 mM Hepes pH 7.5, 50 mM KCl, 1 mM EGTA, 3 mM MgCl_2_, 1 mM DTT, 0.1 mM ATP) overnight at 4°C and loaded onto a 2-ml Glutathione-Sepharose 4B (GE Healthcare Life Sciences, Piscataway, NJ, USA) column pre-charged with GST-N-WASp-VCA (Nolen & Pollard, 2008; Doolittle *et al*, 2013, 3; Antkowiak *et al*, 2019). The column was washed with HKME buffer and bound Arp2/3 was eluted with 20 mM Tris-HCl pH 7.5, 25 mM KCl, 200 mM MgCl_2_, 1 mM EGTA and 1 mM DTT. The presence of protein was detected by using the Bradford reagant, fractions containing protein were pooled, concentrated with an Amicon Ultra 4-ml device (Merck4Biosciences, Darmstadt, Germany), and dialyzed against HKG buffer. Concentrated Arp2/3 was flash frozen in liquid nitrogen and kept at –80°C.

#### WASp (Las17), Capping Protein, ADF/cofilin and Profilin

Rosetta 2(DE3)pLysS cells were transformed with a plasmid designed for *S. cerevisiae* Las17 (Gst-Las17(375-Cter)-6xHis) overexpression. Bacterial cells were collected by centrifugation and then lysed in 20 mM Tris-HCl, pH 7.5, 1 mM DTT, 1 mM EDTA, 200 mM NaCl, 0.1% Triton X-100, 5% glycerol and protease inhibitors (Complete Protease Inhibitor Cocktail, Roche). The lysate was centrifuged at 160,000 x g for 20 min, the supernatant incubated with Glutathione-Sepharose beads, and the protein was purified from the extract. Bound proteins were then eluted with 100 mM L-glutathione reduced and subjected to a second purification by addition of Nickel-Sepharose beads 6 Fast Flow (GE Healthcare Life Sciences, Piscataway, NJ, USA). The protein was eluted with HKI500 buffer (20 mM Hepes, pH 7.5, 200 mM KCl, 500 mM imidazole, pH 7.5), concentrated with an Amicon Ultra 4-ml device and dialyzed against HKG buffer. Protein was flash frozen in liquid nitrogen and kept at –80°C.

*S. cerevisiae* capping protein, ADF/cofilin and profilin were purified as in (Gressin *et al*, 2015). Briefly, proteins were overexpressed in Rosetta 2(DE3)pLysS cells. Cultures were lysed, centrifuged and supernatant were incubated with Nickel-Sepharose beads 6 Fast Flow in HKI20 buffer (20mM Hepes pH 7.5, 200 mM KCl, 20 mM imidazole pH 7.5, 0,1% Triton X-100, 10% glycerol). Proteins were eluted with HKI500 buffer and dyalized against HKG buffer. They were then flash frozen in liquid nitrogen and kept at –80°C.

The labeling of ADF/cofilin was performed using an ADF/cofilin D34C mutant (Gressin *et al*, 2015). Yeast ADF/cofilin D34C mutant was bound to Nickel-Sepharose beads 6 Fast Flow as described above for the wild-type protein. A 5 fold-excess of Alexa Fluor 488 C5-maleimide (Thermo Fisher Scientific) was added overnight at 4°C. Bound protein was cleared from unbound fluorophore before elution in HKI500 buffer, dialyzed against HKG buffer, flash frozen and kept at –80°C.

#### Tropomyosin

Rosetta 2(DE3)pLysS cells were transformed with a plasmid designed for *S. cerevisiae* tropomyosin Tpm1p overexpression. This tropomyosin was modified to contain an Ala-Ser extension at the N-terminal, which mimics its acetylation, and was purified based on a protocol modified from (Skau *et al*, 2009). Briefly, bacteria overexpressing tropomyosin were lysed by sonication in a buffer (50 mM imidazole-HCl, pH 6.9, 300 mM KCl, 5 mM MgCl_2_, 0.3 mM phenylmethylsulfonyl fluoride) supplemented with protease inhibitors (Complete Protease Inhibitor Cocktail, Roche)). Cells were then boiled for 10 min, and the resulting mixture was centrifugated at 300,000 x g for 20 min. The supernatant which contains pure tropomyosin was dialyzed overnight at 4°C against a dialysis buffer (50 mM KCl, 10 mM Tris-HCl, pH 7.5 and 0.5 mM DTT). Tpm1p labeling was performed with the same strategy used for fission yeast tropomyosin Cdc8 labeling (Christensen *et al*, 2017). We mutated *tpm1*’s histidine 114 by site directed mutagenesis to introduce a cysteine (H114C). Immediately after tropomyosin Ala-Ser-Tpm1p H114C purification, the protein was labeled by incubation with a 5-fold excess Alexa Fluor 488 C5-maleimide over tropomyosin overnight at 4°C, and separated on a Sephadex G-25 gel filtration column. The purified fluorescent protein was flash frozen in liquid nitrogen and kept at –80°C.

### Branched- and linear-actin network assembly from microbeads

#### Functionalization of beads

Polystyrene microspheres (2 μm diameter, 2.5% solid (w/v) aqueous suspension, Polysciences, Inc) were washed with HK buffer (20 mM Hepes pH 7.5, 150 mM KCl), diluted 10 times and incubated with 1 μM Las17 for 30 min on ice. Beads were saturated with 1% bovine serum albumin (BSA) for 15 min, washed and stored on ice in HK buffer supplemented with 0.1% BSA. Similarly, glutathione-coated particles (4.37 μm diameter, 0.5% solid (w/v) aqueous suspension, Spherotech, Inc) were coated with GST-Bni1 (1 μM) and then saturated with 1% BSA, washed and stored in HK 0.1% BSA.

#### Branched and linear network reconstitution

Unlabeled and fluorescent actins were mixed to reach a final concentration of 40 μM and a labeling percentage of 1%. Actin polymerization was induced by the addition of G-Buffer and 1x KMEI (50 mM KCl, 1 mM MgCl_2_, 1 mM EGTA, 10 mM imidazole Fluorescence Blank pH 7.8) for 1 h at RT. Las17- and Bni1-coated beads were incubated with F-actin and a minimal set of proteins in a motility buffer (50 mM KCl, 5 mM Hepes, 2.4 mM MgCl_2_; 4 mM DTT; 1 mM ATP; 0.36% methylcellulose 1500 cP and 1.5% BSA) which triggers actin assembly. Standard optimal protein concentrations were 8 μM F-actin, 15 μM profilin, 1 μM capping protein, 500 nM Arp2/3 complex and 600 nM ADF/cofilin. When fluorescent proteins were used, their concentrations were 600 nM for Alexa 488-ADF/cofilin (in which case no black ADF/cofilin was added) and 1 μM for Alexa 488-tropomyosin.

#### Image acquisition, processing and analysis

Images of several beads were acquired 30 min. after the initiation of the experiment on a Zeiss Axio Observer Z1 equipped with a 100x/1.4NA Oil Ph3 Plan-Apochromat objective and a Hamamatsu ORCA-Flash 4.0LT camera. Images were acquired with Zen 2.3 blue edition using the same light intensity and exposure time.

#### Data quantification for biomimetic assays

Fluorescence intensity of actin networks and fluorescent ABPs was quantified using Fiji (Version 1.52p), and fluorescence of the background was substracted. Similarly to the *in vivo* actin network deviation index, a linear-to-branched ratio was calculated to compare the efficiency of actin assembly *in vitro* for both branched- and linear-networks of actin filaments for a given biochemical condition. This index measures how actin assembly between branched and linear networks deviates from the values obtained when *S. cerevisiae* actin is used. It is defined as follows:

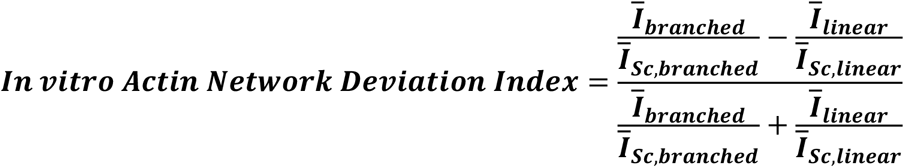

Where 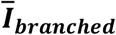 is the average normalized intensity of the branched network for all Las17 beads with a given actin, 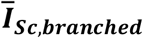 is the same value for *S. cerevisiae* actin, 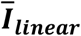 is the average normalized intensity of the linear network for all Bni beads with a given actin and 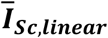 is the same value for *S. cerevisiae* actin.

### Actin-ABP contact analysis

The amino acid positions of the substitutions between *Saccharomyces cerevisiae* (Act_Sc), Node 2 (Act_N2), and *Candida albicans* (Act_Ca) actin sequences were inspected within high resolution X-ray crystal and cryoEM structures of complexes containing actin. The PDB accession codes are: G-actin (1YAG) (Vorobiev *et al*, 2003), F-actin (6DJN) (Chou & Pollard, 2019), ADF/cofilin (5YU8 and 1CFY) (Tanaka *et al*, 2018; Fedorov *et al*, 1997)), Arp2/3 daughter filament (6W17) (Shaaban *et al*, 2020), profilin (1YPR and 3CHW) (Baek *et al*, 2008; Eads *et al*, 1998); CP/Arp1 (5ADX) (Urnavicius *et al*, 2015), WH2 (5YPU), formin (1Y64 and 4EAH) (Otomo *et al*, 2005; Thompson *et al*, 2013) and tropomyosin (5JLF) (von der Ecken *et al*, 2016). Actin residues that are within 5 Å of the binding protein were identified in the CCP4 program CONTACT (Winn *et al*, 2011), with the exception of the Arp2/3 mother filament which were within 10 Å since the coordinates were not released when this study was performed (Fäßler *et al*, 2020). All contacts were visually inspected in COOT (Emsley *et al*, 2010).

## Data availability

This study includes no data deposited in external repositories.

## Data reproducibility

All experiments were repeated at least two times. In all plots, error bars indicate standard deviations. As standard deviations were not similar for all experiments, we used Brown-Forsythe and Welch ANOVA tests, with Dunnett’s T3 multiple comparisons tests. P value style: * <0.05, ** <0.01, *** <0.001. Correlations were computed with a two-tailed p value and a confidence interval of 95%, correlation coefficients r correspond to Pearson correlation coefficients.

For the colony area measurements, colonies were measured from 2 plates, results were normalized to control and pulled together. More than 10 colonies per strain were quantified. For western blot measurements, 2 independent samples per strain were loaded twice each and analyzed. For phalloidin stained cells, 30 cells were measured per strain. For the *in vitro* polymerization assays, at least 14 beads were measured from two independent experiments, and data presented in the manuscript correspond to the two sets of experiments pulled together.

## Acknowledgements

The authors thank Isabelle Sagot and Emilia Mauriello for their invaluable advice on the project, and Sarah Würbel for her technical help. This project has received funding from the European Research Council (ERC) under the European Union’s Horizon 2020 research and innovation programme (grant agreement n° 638376/Segregactin) to A.M., from the Labex INFORM (ANR-11-LABX-0054, funded by the ‘Investissements d’Avenir French Government program’), and from the Fondation pour la Recherche Médicale (FRM) to M.B.S. under the program Fin de these (ref. FDT201904008021). R.C.R. thanks Vidyasirimedhi Institute of Science and Technology (VISTEC), RIIS and JSPS (KAKENHI grant number JP20H00476) for support. We acknowledge the France-BioImaging infrastructure supported by the French National Research Agency (ANR-10-INSB-04-01).

## Author contributions

Conceptualization, Methodology and Writing: M.B.S., C.P.T, R.C.R. and A.M. Investigation: M.B.S., C.P.T., A.A., A.G., R.C.R. and A.M. Validation and Visualization: M.B.S. Funding acquisition, Supervision and Project administration: R.C.R. and A.M.

## Abbreviations

FH1-FH2: Formin Homology domain 1-Formin Homology domain 2

## Supplementary Figures legends

**Figure S1, related to Figure 1. Selection of actins strategy. (A)** Complete phylogenetic tree that was used as input for FastML ancestral reconstruction analysis (Ashkenazy *et al*, 2012) **(B)** Posterior probability for the ancestral sequences used in this study, showing high confidence in the predicted sequences. **(C)** (Top) Multiple sequence alignment for all actin sequences used in this study. (Bottom) Schematic representations of actin 3D structure (1YAG, (Vorobiev *et al*, 2003)), with position of amino acid differences shown with colored dots for each actin. **(D)** Schematic representation of mutagenesis strategy by homologous recombination used in this study (see also Methods).

**Figure S2, related to Figure 2. Effect of removing *S. cerevisiae*’s Act1 intron and of silent mutations in the actin gene. (A)** 3-fold serial dilutions of different yeast strains cultures grown at 25°C for 2 days on a YPD plate. **(B)** Quantification of (A) by measurement of colony area. **(C)** Actin expression levels shown by western blotting, with tubulin (Tub1p) as a loading control. **(D)** Quantification of actin expression levels. **(E)** Phallodin stain depicting F-actin organization. Images are maximum intensity projections of 3D stacks. Scalebar: 3 μm. **(F)** *In vivo* actin network deviation indexes. **(G)** Polarity indexes. **(H)** Multiple sequence alignment of the beginning of the nucleotide sequence (top) and the beginning of the amino acid sequence (bottom), as an example of how we used coding sequences from other organisms that we modified minimally so that the final product remained *S. cerevisiae* actin. **(I)** Actin expression levels as a function of nucleotide conservation, showing that increased number of silent mutations lowers actin expression. **(J)** Colony area as a function of nucleotide identity, showing a threshold of nucleotide conservation (78%<id<82%) below which growth rates drastically reduce. Abbreviations: Sc - wild-type *S. cerevisiae* cells, ScI - *S. cerevisiae* cells where the actin gene has been replaced with the full construct carrying the wild-type gene, ScNI - *S. cerevisiae* cells where the actin gene has been replaced with the wild-type gene but without the intron.

**Figure S3. C4 actin antibody has a higher affinity for rabbit muscle actin than for *S. cerevisiae* actin. (A)** The binding site of the C4 antibody, indicated as “C4_Epitope”, is found on Act_Hs and on rabbit muscle actin. In all other actin variants used in this study, the sequence varies of one amino acid (called here “Mutated_Epitope”) but is recognized by C4 antibody. **(B)** Western blot with equivalent amounts of purified yeast actin and rabbit actin. The amount of protein was revealed by two methods: Ponceau staining and chemiluminescence. The chemiluminescence signal corresponds to the one produced by the secondary antibody after incubation with a primary antibody anti-actin C4 and a secondary antibody conjugated with HRP. **(C)** Quantification of (B) indicates that immunolabeling of rabbit muscle actin with C4 antibody leads to a 1.48-fold more intense signal than immunolabeling of *S. cerevisiae* actin.

## Supplementary Tables legends

**Table S1. Complete list of all actins used for the ancestral sequence reconstruction using FastML.**

**Table S2. List of plasmids used in this study.** All plasmids were done in a pGEX-4T1 backbone.

**Table S3. List of yeast strains in this study.**

